# Inference of epistatic effects leading to entrenchment and drug resistance in HIV-1 protease

**DOI:** 10.1101/063750

**Authors:** William F. Flynn, Allan Haldane, Bruce E. Torbett, Ronald M. Levy

## Abstract

Understanding the complex mutation patterns that give rise to drug resistant viral strains provides a foundation for developing more effective treatment strategies for HIV/AIDS. Multiple sequence alignments of drug-experienced HIV-1 protease sequences contain networks of many pair correlations which can be used to build a (Potts) Hamiltonian model of these mutation patterns. Using this Hamiltonian model we translate HIV protease sequence covariation data into quantitative predictions for the probability of observing specific mutation patterns which are in agreement with the observed sequence statistics. We find that the statistical energies of the Potts model are correlated with the fitness of individual proteins containing therapy-associated mutations as estimated by in vitro measurements of protein stability and viral infectivity. We show that the penalty for acquiring primary resistance mutations depends on the epistatic interactions with the sequence background. Primary mutations which lead to drug resistance can become highly advantageous (or entrenched) by the complex mutation patterns which arise in response to drug therapy despite being destabilizing in the wildtype background. Anticipating epistatic effects is important for the design of future protease inhibitor therapies.

## I. INTRODUCTION

The ability of HIV to rapidly mutate leads to antiretroviral therapy (ART) failure among infected patients. Enzymes coded by the *pol* gene play critical roles in viral maturation and have been key targets of several families of drugs used in combination therapies. The protease enzyme is responsible for the cleavage of the Gag and Gag-Pol polyproteins into functional constituent proteins and it has been estimated that resistance develops in as many as 50% of patients undergoing monotherapy (Richman et al. 2004) and as many as 30% of patients undergoing modern combination antiretroviral therapy (c-ART) (Gupta et al. 2008).

The combined selective pressures of the human immune response and antiretroviral therapies greatly affect the evolution of targeted portions of the HIV-1 genome and give rise to patterns of correlated amino acid substitutions. As an enzyme responsible for the maturation of the virion, the mutational landscape of HIV protease is further constrained due to function, structure, thermodynamics, and kinetics (Bloom et al. 2010, Haq et al. 2012, Lockless et al. 1999, Zeldovich and Shakhnovich 2008, Zeldovich et al. 2007). As a consequence of these constraints, complex mutational patterns often arise in patients who have failed c-ART therapies containing protease inhibitors (PI), with mutations located both at critical residue positions in or near the protease active site and others distal from the active site (Chang and Torbett 2011, Flynn et al. 2015, Fun et al. 2012, Haq et al. 2012). In particular, the selective pressure of PI therapy gives rise to patterns of strongly correlated mutations generally not observed in the absence of c-ART, and more therapy-associated mutations accumulate under PI therapy than under all other types of ART therapies (Shafer 2006, Shafer and Schapiro 2008, Wu et al. 2003). In fact, the majority of drug-experienced subtype B protease sequences in the Stanford HIV Drug Resistance Database (HIVDB) have more than 4 PI-therapy-associated mutations (see Figure S1). Within the Stanford HIVDB are patterns of multiple resistance mutations, and in order to overcome the development of resistance, understanding these patterns is critical.

A mutation’s impact on protein stability or fitness depends on the genetic background in which it is acquired. Geneticists call this phenomenon “epistasis”. It is well understood that major drug resistance mutations in HIV protease destabilize the protease in some way, reducing protein stability or enzymatic activity, which can greatly alter the replicative and transmissive ability, or *fitness*, of that viral strain (Bloom et al. 2010, Boucher et al. 2016, Grenfell et al. 2004, Wang et al. 2002). To compensate for this fitness loss, protease accumulates accessory mutations which have been shown to restore stability or activity (Chang and Torbett 2011, Fun et al. 2012, Martinez-Picado et al. 1999). But it is unclear how the acquisition and impact of primary and accessory mutations are modulated in the presence of the many different genetic backgrounds observed, especially those present in the complex resistant genotypes that arise under inhibitor therapy.

Coevolutionary information derived from large collections of related protein sequences can be used to build models of protein structure and fitness (Burger and van Nimwegen 2010, Göbel et al. 1994, Hinkley et al. 2011, Liu et al. 2009, Lockless et al. 1999, Socolich et al. 2005). Given a multiple sequence alignment (MSA) of related protein sequences, a probabilistic model of the network of interacting protein residues can be inferred from the pair correlations encoded in the MSA. Recently, probabilistic models, called Potts models, have been used to assign scores to individual protein sequences which correlate with experimental measures of fitness (Ferguson et al. 2013, Figliuzzi et al. 2015, Haq et al. 2012, Hopf et al. 2015, Mann et al. 2014). These advances build upon previous and ongoing work in which Potts models have been used to extract information from sequence data regarding tertiary and quaternary structure of protein families (Barton et al. 2016a, Haldane et al. 2016, Jacquin et al. 2016, Marks et al. 2012, Morcos et al. 2011, 2014, Sulkowska et al. 2012, Sutto et al. 2015, Weigt et al. 2009) and sequence-specific quantitative predictions of viral protein stability and fitness (Barton et al. 2016b, Butler et al. 2016, Haq et al. 2012, Shekhar et al. 2013).

In this study, we show how such models can be constructed to capture the epistatic interactions involved in the evolution of drug resistance in HIV-1 protease. The acquisition of resistance mutations which accumulate under the selective pressure of inhibitor therapy leave many residual correlations observable in MSAs of drug-experienced sequences (Hoffman et al. 2003, Rhee et al. 2007, Wu et al. 2003), and we use the pair correlations that can be extracted from MSAs to construct a Potts model of the mutational landscape of drug experienced HIV-1 protease. We first provide several tests which demonstrate that our inferred model faithfully reproduces several key features of our original MSA including higher order correlations. We then compare the Potts model statistical energies with experimental measurements of fitness, including structural stability and relative infectivity of individual HIV protease variants which contain resistance mutations. Finally, the Potts scores are used to describe the epistatic mutational landscape of three primary resistance mutations. We observe strong epistatic effects. The primary mutations are destabilizing in the context of the wildtype background, but become stabilizing on average as other resistance mutations accumulate in the background, similar to the concept of entrenchment in systems biology (Gong et al. 2013, Pollock et al. 2012, Shah et al. 2015). Furthermore, we find that entrenchment is modulated by the collective effect of the entire sequence, including mutations at polymorphic residues, and the variance of the statistical energy cost of introducing a primary mutation increases as resistance mutations accumulate; this heterogeneity is another manifestation of epistasis (Barton et al. 2016b, McCandlish et al. 2015). These findings provide a framework for exploring mutational resistance mechanisms using probabilistic models.

## II. RESULTS

### A. Model inference and dataset

Given a multiple sequence alignment (MSA), we can infer a statistical model 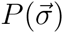 for the probability of finding a protein sequence with sequence identity 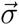 which takes the form 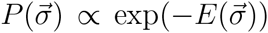 from the statistical properties of the MSA. The maximum entropy model which reproduces the first and second order marginal distributions of the MSA, *P*_*i*_(*σ*_*i*_) and *P*_*ij*_(*σ*_*i*_, *σ*_*j*_) of residue positions *i* and position pairs *i*, *j*, is given by the Potts Hamiltonian 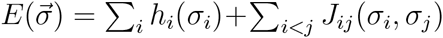, where the fields *h*_*i*_(*σ*_*i*_) and couplings *J*_*i,j*_(*σ*_*i*_, *σ*_*j*_) represent the preference for residue *σ*_*i*_ at position *i* and residue pair *σ*_*i*_*σ*_*j*_ at positions *i*, *j*, respectively. The Potts model is fit to the bivariate marginals of the MSA such that it recovers the correlated pair information *C*_*ij*_(*σ*_*i*_, *σ*_*j*_) = *P*_*ij*_(*σ*_*i*_, *σ*_*j*_) − *P*_*i*_(*σ*_*i*_)*P*_*j*_(*σ*_*j*_).

The Potts model captures epistatic effects; in contrast an independent model of a multiple sequence alignment can be constructed by summing the logarithm of the univariate marginals 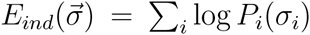. As described in the following section, the ability to reproduce higher order marginals of the MSA (beyond second order) is a true predictive test of the Potts model, one which the independent model fails.

As described in the introduction, protease sequence evolution under protease inhibitor selective pressure produces correlations between amino acid substitutions that are larger in magnitude than those that occur in the absence of drug pressure (seen in Figure S2) (Gupta and Adami 2016, Rhee et al. 2007, Wu et al. 2003). Although correlations between some drug-associated sites can be identified through analysis of drug-naive sequences, or structural and/or evolutionary constraints (Butler et al. 2016, Hoffman et al. 2003), the most complete model of the epistatic landscape of drug-resistance mutations is constructed using the correlations found in a varied set of drug-experienced sequences. As we demonstrate in later sections, correlations among the primary, accessory, and polymorphic mutations which arise under c-ART therapy all contribute to protease fitness. Starting with an MSA constructed from 5610 HIV-1 subtype B drug-experienced protease sequences obtained from the Stanford HIVDB, we have inferred a Potts model using a Markov Chain Monte Carlo (MCMC) method implemented on GPUs (see Materials and Methods and the supplemental information of Haldane et al. (Haldane et al. 2016) for more details).

### B. Recovery of the observed sequence statistics – marginal probabilities

We can gauge the accuracy of the model by examining how well the model reproduces various statistics of the MSA. The most direct test is the reproduction of higher order correlations observed in the multiple sequence alignment beyond pair correlations. Shown in Figure 1A is the recovery of the marginal probabilities of the most common subsequences observed in the dataset across varying subsequence lengths. The recovery of the bivariate marginals (pair frequencies) is not predictive but it demonstrates the quality of fit of the Potts model. The results shown in Figure 1 demonstrate that the Potts model is able to predict the frequencies of higher order marginals with accuracy. The Pearson correlation coefficient for the observed probabilities compared with the Potts model prediction remains above *R*^2^ ≥ 0.95 for subsequence lengths as large as 14. In contrast the independent model correlation coefficient is significantly worse (*R*^2^ → 0.22).

**FIG. 1:**
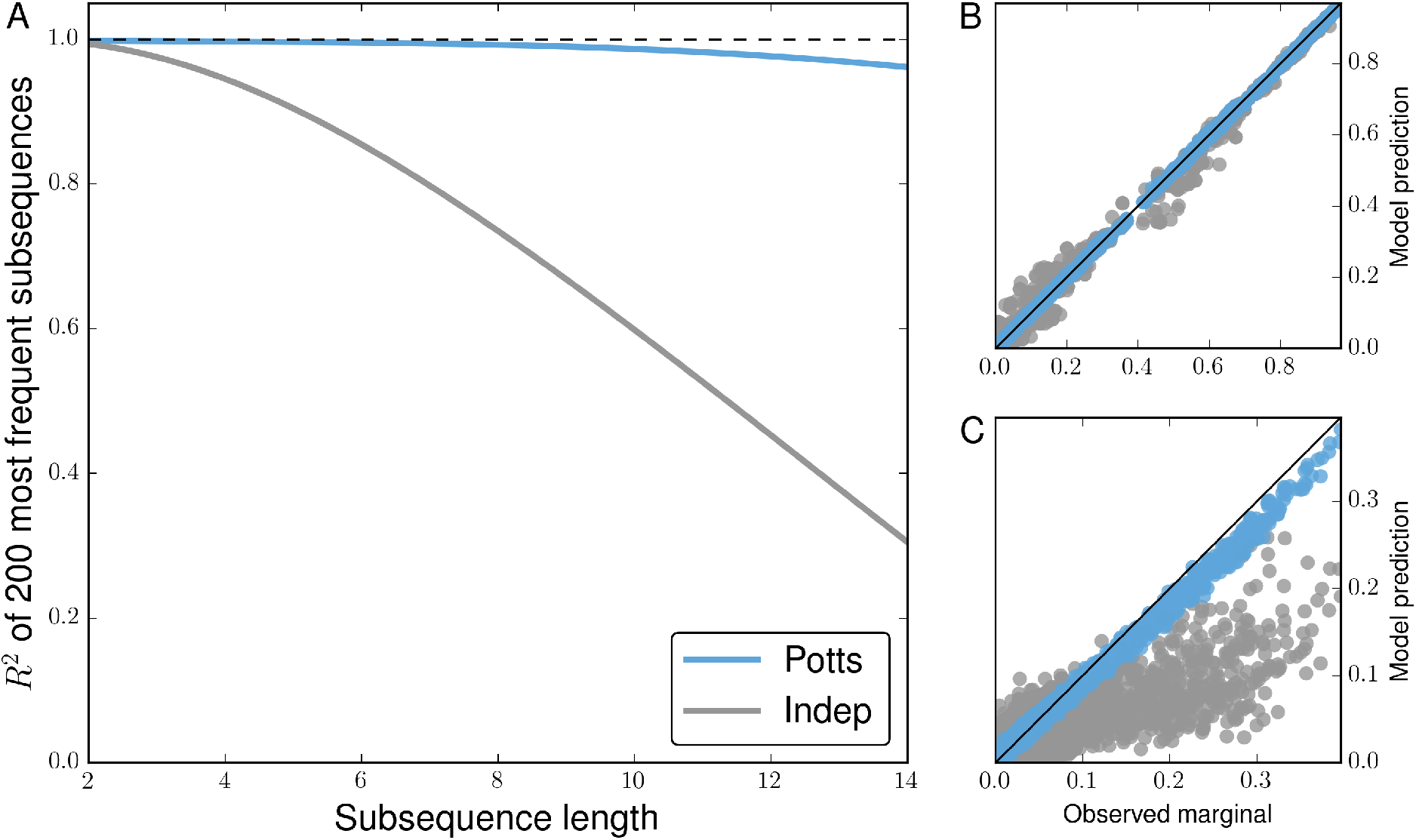
Potts model is predictive of higher order sequence statistics. For each subsequence length varying from 2 to 14, subsequence frequencies determined by counting occurences in the MSA are computed for all observed subsequences at 500 randomly chosen combinations among 36 PI-associated positions. (A) Pearson R^2^ of the 200 most probable observed subsequence frequencies (marginals) with corresponding predictions by Potts (blue) and independent (gray) models for varying subsequence lengths. (B) 2^nd^ and (C) 14^th^ order observed marginals predicted by both models. Shown in (B,C) are observed frequencies at the 500 randomly chosen combinations of 2 and 14 positions among 36 PI-associated sites, with approximately 2500 and 5600 subsequence frequencies greater than 0.01 visible, respectively.

Figure 2 shows the probability distribution of sequences that differ from the consensus by *k* mutations as predicted by the Potts and independent models compared to the observed distribution derived from the MSA. The Potts model predicts a distribution of mutations per sequence which is very close to the observed distribution whereas the independent model incorrectly predicts a multinomial distribution centered about 8 mutations from consensus. The very good agreement between the higher order sequence statistics of the Potts model and the observed statistics from the MSA provides additional evidence that the Potts model is not overfit.

**FIG. 2:**
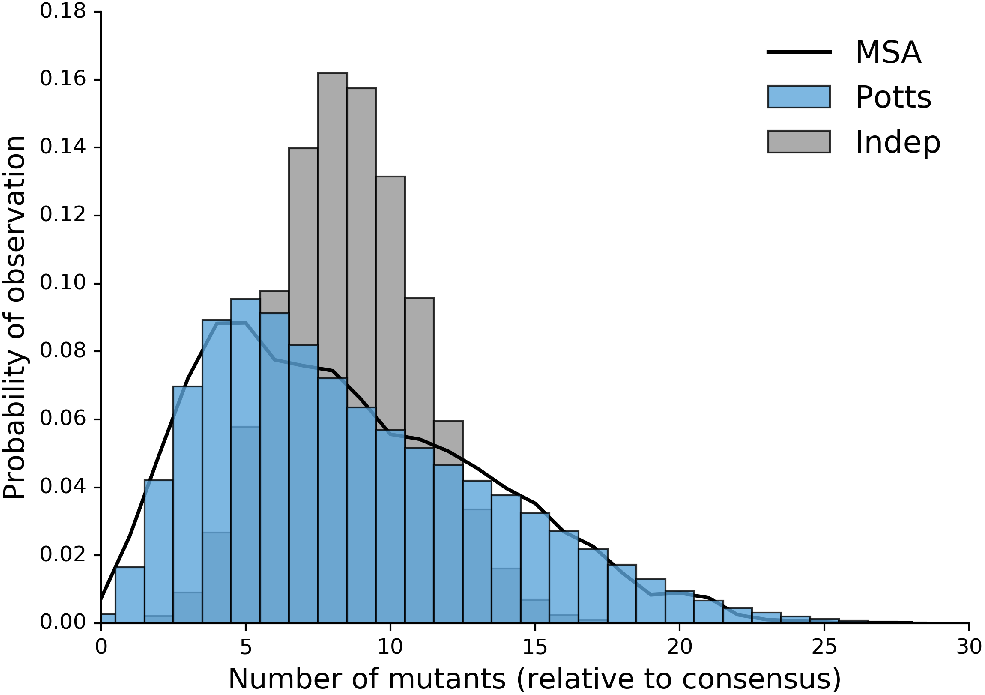
Potts model captures properties of full length sequence ensemble. Probabilities of observing sequences with any *k* mutations relative to the consensus sequence as observed in original MSA (black) and predicted by the Potts (blue) and independent (gray) models.

The Potts model also captures the observed statistics for larger subsequences, but as subsequence lengths increase, observed marginal probabilities in our MSA approach the sampling limit of the alignment (1/*N* ≈ 2 × 10^−4^), meaning comparisons with the observed data at this level become dominated by noise. Tests with synthetic data (not shown) confirm that for longer subsequences the discrepancy between observed higher order marginals and the Potts model are consistent with effects caused by the finite sample size (5610 sequences) of the MSA (Haldane et al. 2016). In the following section, we compare Potts model statistical energies with experimentally determined measurements of protease fitness.

### C. Protease mutations, protein stability, and replicative capacity

Two experimental tests used to quantify the effects of protease mutations on viral fitness are thermal stability of the folded protein and replicative capacity (Chang and Torbett 2011, Louis et al. 2011, Muzammil et al. 2003). Chang and Torbett demonstrate that stability is compromised by the acquisition of primary mutations and this loss of stability can be rescued by known compensatory mutations, sometimes in excess of the reference stability. Muzammil et al. and Louis et al. have shown that patterns of up to 10 or more resistance mutations do not necessarily suffer from reduced stability relative to the wildtype, and that non-active site mutations can lead to resistance in certain sequence contexts. In Figure 3A the change in statistical Potts energies, Δ*E* = *E* − *E*_*ref*_ is plotted versus the change in thermal stability, where *E* and *E*_*ref*_ are the statistical energies of the mutated and reference sequences corresponding to each pair of stability measurements. We observe a strong correlation between Potts Δ*E* and the change in stability as reflected by the change in melting temperature (*R* = −0.85, *p* = 0.0003). In contrast, the change in stability computed using the independent model shows no correlation (Figure S3A).

**FIG. 3:**
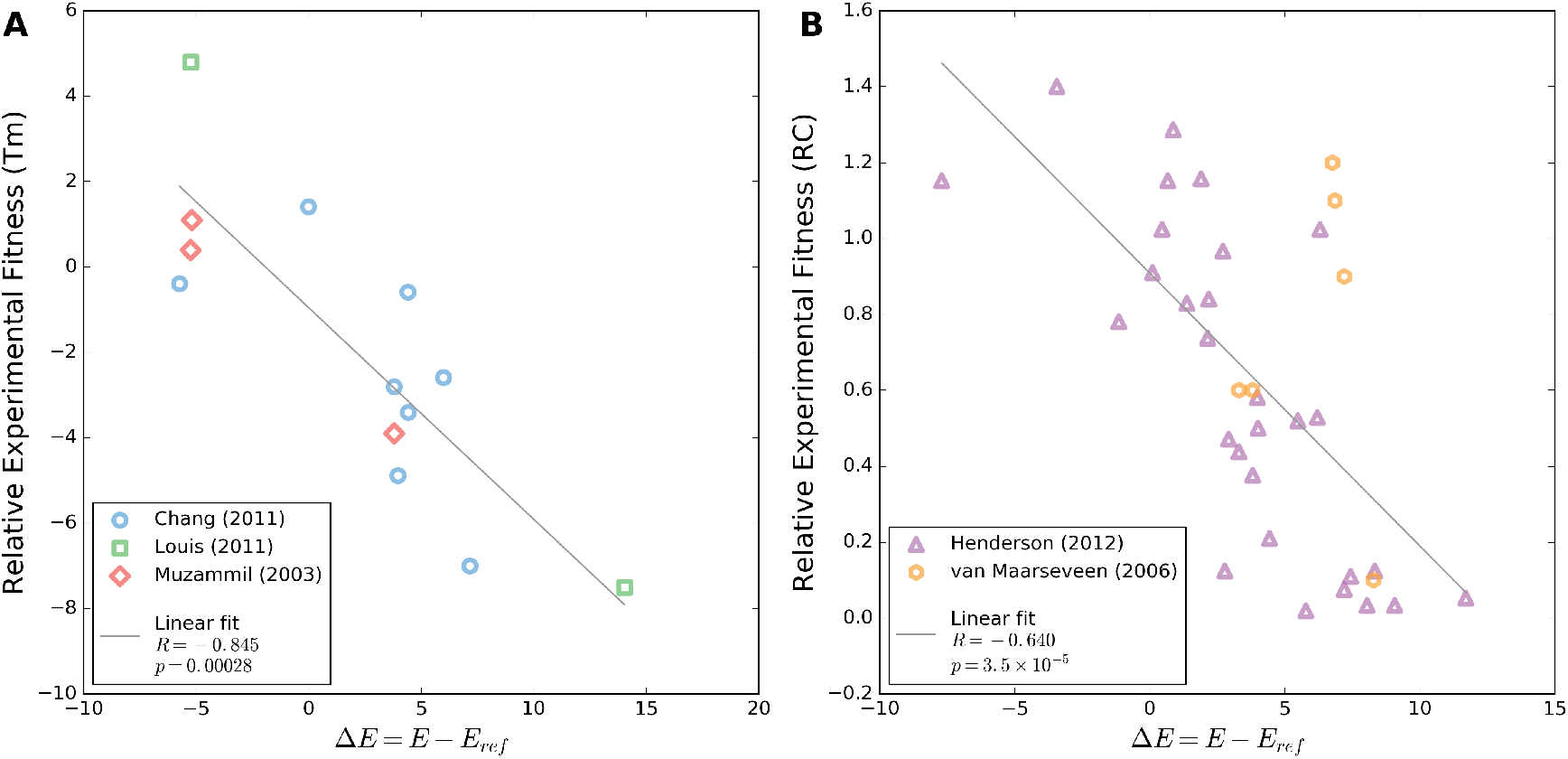
Change in Potts energy correlates with change in experimental fitness. (A) Changes in melting temperature (*T*_*m*_) for individual sequences relative to a reference sequence extracted from literature (Chang and Torbett 2011, Louis et al. 2011, Muzammil et al. 2003). These sequences differ from the wildtype by 1–2 mutations (Chang and Torbett 2011) up to 10–14 mutations (Louis et al. 2011, Muzammil et al. 2003). (B) Change in relative infectivty as measured by replicative capacity assay for individual sequences containing only single point mutations (Henderson et al. 2012) and 1–5 mutations (van Maarseveen et al. 2006). In both panels a linear regression fit with Pearson’s R and associated two-tailed p-value are provided in the legend.

We have extracted results for viral replicative capacity in which 29 single Protease mutants were studied by Henderson et al. (Henderson et al. 2012) and an additional small set of more complex sequence variants (van Maarseveen et al. 2006) that were tested relative to the wildtype sequence. As with the stability measurements, we find the relative Potts energy correlates well with infectivity (*r* = −0.64, *p* < 10^−5^), shown in Figure 3B. The same comparison using the independent model computed fitness again shows no predictive power (Figure S3B). Complementary to the RC assay presented in their study, Henderson et al. presented a SpIn assay and an additional assay measuring drug concentrations which inhibit protease function (EC50). Potts fitness predictions against these data are shown in Figure S4. While this additional comparison does not show statistically significant correlation, probably because the observed measurements span a much smaller range of values, they do exhibit the same negative trends observed in Figure 3. All data shown in Figures 3, S3, S4 can be found in Supplementary Data 1.

The results presented here are reinforced by other recent studies of protein evolutionary landscapes (Ferguson et al. 2013, Figliuzzi et al. 2015, Hopf et al. 2015, Mann et al. 2014) where varying measures of experimental fitness are compared to statistical energies derived from correlated Potts models constructed from multiple sequence alignments. The range of statistical energies and the correlation with fitness are qualitatively similar to those presented by Ferguson et al. and Mann et al. where statistical energies of engineered HIV-1 Gag variants generated using a similar inference technique are compared with replicative fitness assays. The same can be said for correlations between Potts scores and relative folding free energies of Beta Lactamase TEM-1 presented by Figliuzzi et al.. This collection of studies demonstrate that Potts model statistical energies correlate with the fitness of protein sequences in different contexts, including protein families evolving under weak selection pressure (Figliuzzi et al. 2015, Hopf et al. 2015), viral proteins evolving under immune pressure (Ferguson et al. 2013, Mann et al. 2014), and as presented here, viral proteins evolving under drug pressure.

### D. Inference of epistasis among therapy-associated mutations

The sequences present in the Stanford HIVDB have been deposited at many stages of HIV infection and treatment, showcasing a variety of resistance patterns spanning from wildtype to patterns of more than 15 mutations at PI-associated positions. In this section, we describe how Potts statistical energies can be used to infer epistatic effects on the major HIV protease resistance mutations.

Although all current PIs are competitive active site inhibitors, major resistance mutations can be found both inside and outside of the protease active site; the substrate envelope hypothesis suggests this arises because PIs have a larger interaction surface with protease compared to that of its natural substrates (King et al. 2004, Özen et al. 2011, Prabu-Jeyabalan et al. 2002). V82 and I84 are positions inside the substrate cleft and major resistance mutations V82A and I84V have been shown to directly affect binding of inhibitors (Chellappan et al. 2007, King et al. 2002, Lefebvre and Schiffer 2008). L90 is a residue located outside of the substrate cleft and flap sites. Mutations at position 90, specifically L90M, have been shown to allow shifting of the aspartic acids of the active site catalytic triad (D25) on both chains, subsequently allowing for larger conformational changes at the dimer interface and active site cleft that reduce inhibitor binding (Kovalevsky et al. 2006, Mahalingam et al. 2004, Ode et al. 2006).

Given a sequence containing one of the 3 mutants V82A, I84V, and L90M, we can determine the context-dependence of that mutation in its background by calculating the change in statistical energy associated with reversion of that mutation back to wildtype. This corresponds to computing Δ*E* = *E*_obs_ − *E*_*rev*_ where *E*_*obs*_ is the Potts energy of an observed sequence with one of these primary mutations and *E*_*rev*_ is the Potts energy of that sequence with the primary mutation reverted to its consensus amino acid type. Due to the pairwise nature of the Potts Hamiltonian, this computation reveals a measure of epistasis for a sequence 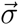 containing mutant *X* → *Y* at position *k*

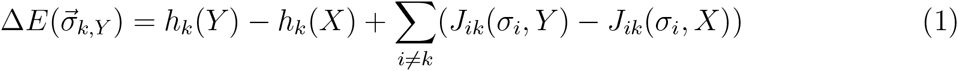

where the pair terms *J*_*ik*_ are the couplings between the mutation site and all other positions in the background. When this measure is positive, the background imparts a fitness penalty for the reversion of the primary resistance mutation to the wildtype and when negative, the sequence regains fitness with reversion to wildtype. Using this measure, we computed Δ*E* for every sequence in our HIVDB MSA containing V82A, I84V, L90M and have arranged the energies versus sequence hamming distance from the consensus including only PI-associated sites, shown in Figure 4. As more mutations accumulate in the background, the preference for each primary resistance mutation to revert to wildtype is lost and the primary mutation becomes preferred over the wildtype on average when enough background mutations have accumulated. These crossover points are 6, 9, and 7 mutations for V82A, I84V, and L90M, respectively. When a sufficient number of mutations have accumulated, the primary resistance mutation becomes *entrenched*, meaning a reversion to wildtype at that position is destabilizing in most sequences; the primary mutation becomes more entrenched as more background mutations are acquired. The effect is largest for L90M; for sequences containing > 7 PI-associated mutations, on average the L90M primary mutation is ≈100 times more likely than the wildtype leucine at position 90. In contrast, this primary mutation is ≈80 times less likely than the wildtype residue in the subtype B consensus sequence background. In other words, there is an ≈8, 000 fold difference in the probability of observing the mutation L90M depending on the background sequence. The trend shared for V82A, I84V, and L90M is representative of the larger class of primary mutations; mutations V32I, M46L, I47V, G48V, I50V, I54V, L76V, and others become less destabilizing as the number of background mutations increases (see Figure S5).

**FIG. 4:**
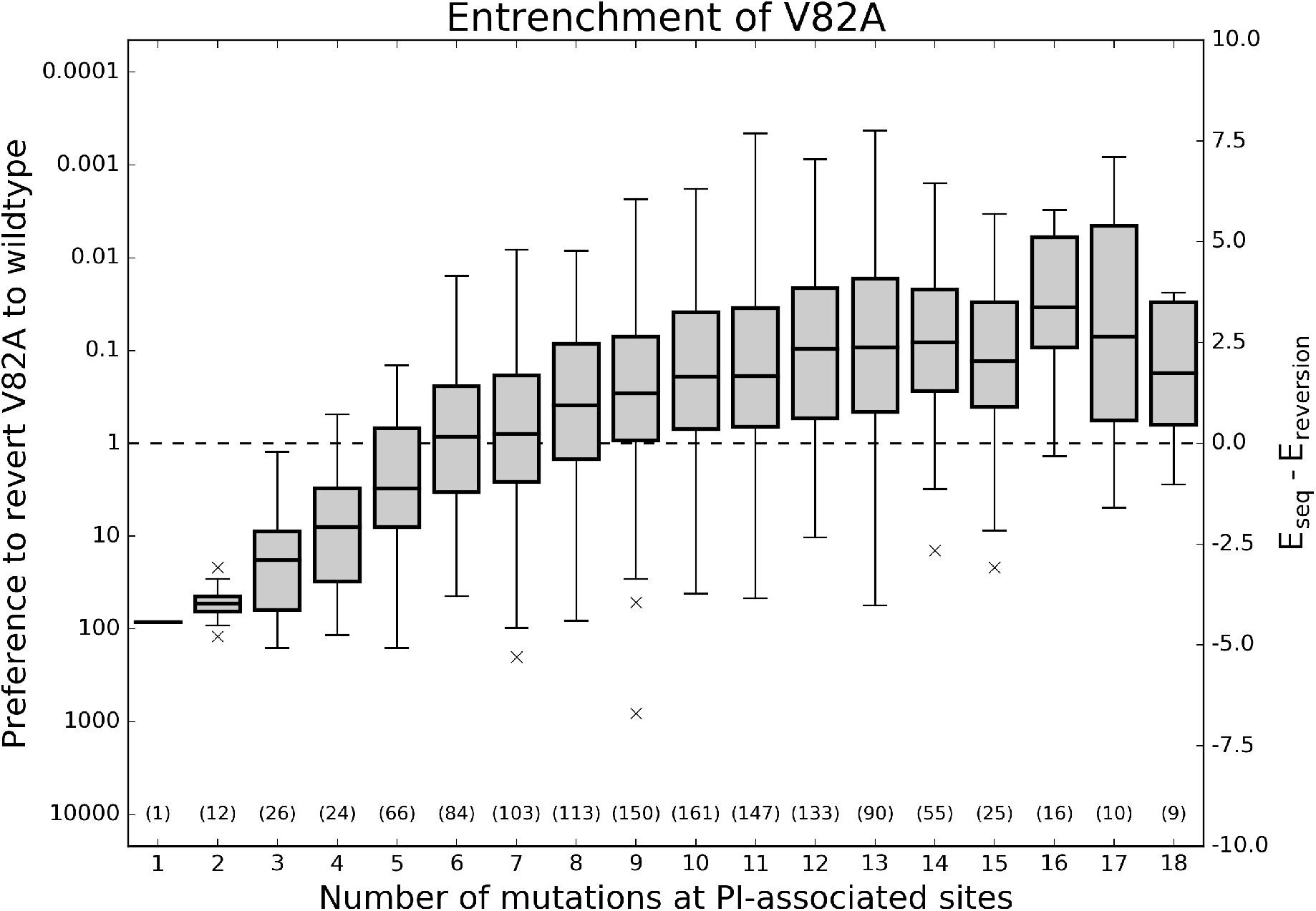
Effect of epistasis on the fitness penalty incurred by primary resistance mutations. For each of the 3 primary HIV protease mutations described in (Chang and Torbett 2011), two Potts statistical energies are computed for all observed sequences containing that mutation: *E*_*seq*_, the energy of the sequence with that mutation and *E*_*reversion*_, the energy with that primary mutation reverted to wildtype. This Potts energy difference, Δ*E* = *E*_*seq*_ − *E*_*reversion*_ is shown versus hamming distance from the wildtype including only PI-assocated positions. Ordinate scales are given in both relative probability of reversion exp(−Δ*E*) (left) and Δ*E* (right). Values below (above) the dashed line on the ordinate correspond to fitness gain (penalty) upon reversion to wildtype. Although primary resistance mutations initially destabilize the protease, as mutations accumulate, the primary resistance mutations become entrenched, meaning their reversion becomes destabilizing to the protein.

Why are primary resistance mutations much more likely in some backgrounds and not others? Are these effects caused by a small set of epistatic interactions with the primary resistance mutation or the collective effect of many small epistatic interactions?

To answer these questions, we compared the sequence backgrounds which most entrench primary mutations from those sequences which most prefer wildtype instead of the primary mutation. Using as an example a fixed hamming distance of 10 from the subtype B consensus sequence, we examined the differences between the sequences among the top 10% and bottom 10% of Δ*E* values in the *h* = 10 column in each of the subplots of Figure 4. The *h* = 10 column was chosen as it is the column with the most data for the primary mutations V82A, I84V, and L90M. These two groups of sequences, top 10% and bottom 10%, are referred to as “most entrenched” (ME) and “least entrenched” (LE) sequences, respectively.

One might expect that the accumulation of accessory mutations in a sequence will lead to the entrenchment of a primary mutation and, under this assumption, the most entrenched sequences should contain more accessory mutations than the least entrenched sequences. We observe more accessory mutations in the most entrenched sequences on average, but the difference is not significant and a large number of accessory mutations accumulate in the least entrenching sequences for V82A, I84V, and L90M as shown in Figure 5. In other words, simply counting accessory mutations in a sequence is unlikely to predict whether that sequence will entrench a primary mutation.

**FIG. 5:**
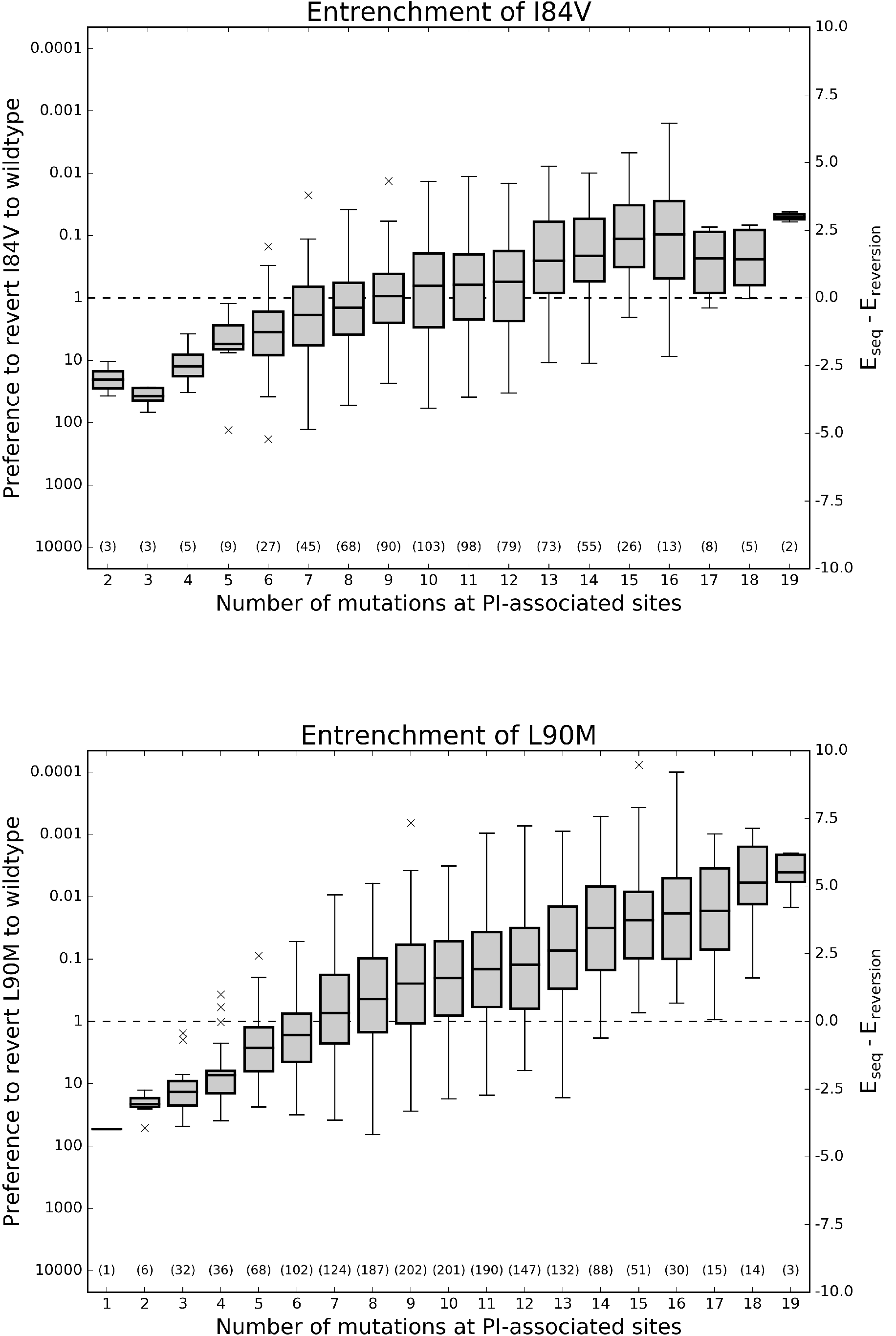
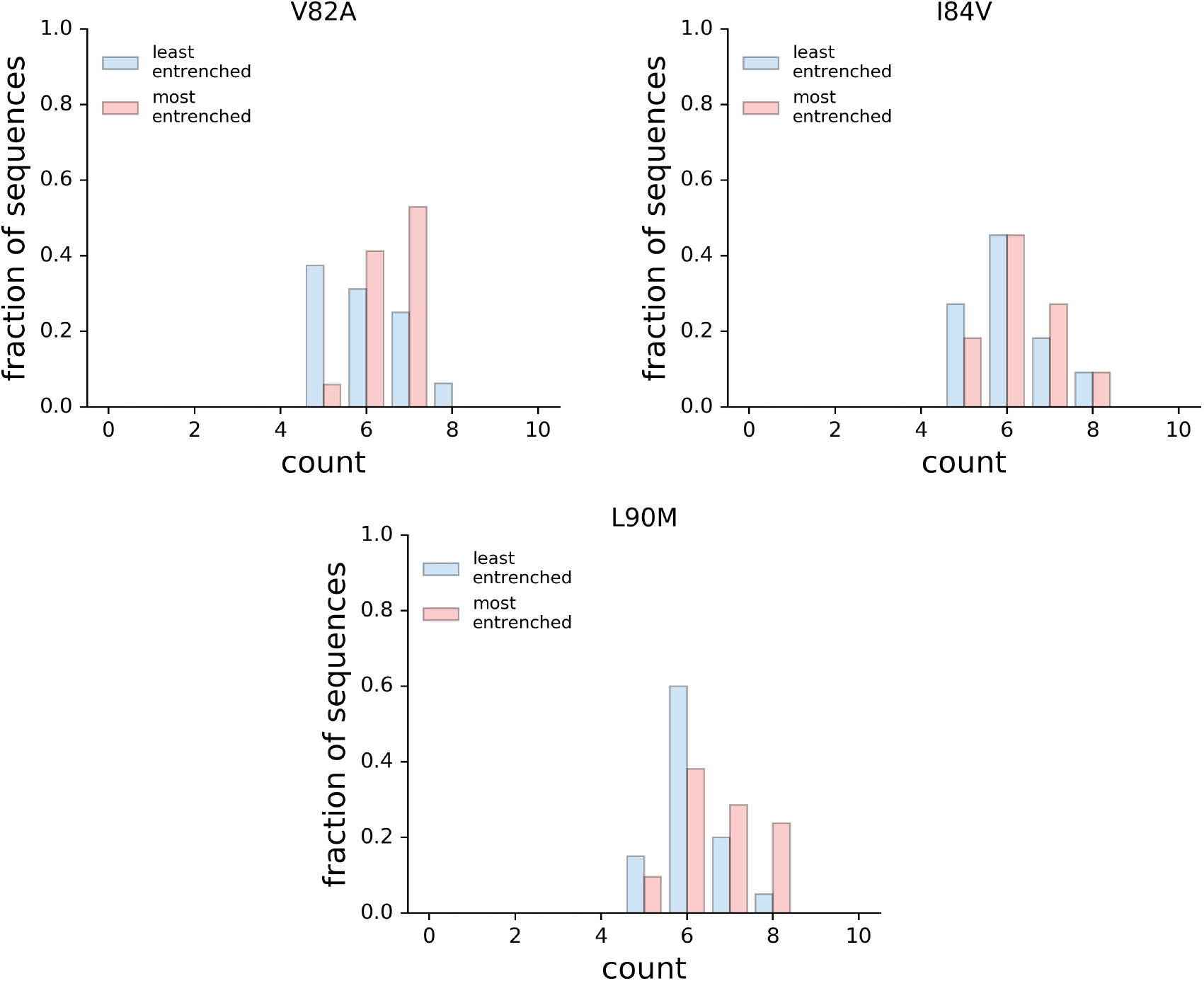
Distributions of accessory mutations in most and least entrenching sequences. The number of accessory mutations among the 10% most and least entrenching sequences for the primary mutations V82A, I84V, and L90M with a fixed hamming distance of 10 from consensus. In all three cases, the distributions are not significantly different (Mann Whintey *U*_*V*82_ = 92.5, *U*_*V*84_ = 53.0, *U*_*L*90_ = 145.5, all with p > 0.05).

Previous research has identified significant correlations between various primary and accessory mutations and the primary resistance mutations under study here (Flynn et al. 2015, Rhee et al. 2007, Wu et al. 2003). We find that the presence of these accessory mutations alone cannot account for the separation of the most entrenched sequences from the least entrenched sequences. The most striking example is the double mutant G73S-L90M. G73S is present in 75% of the ME sequences and never present in the LE sequences; however, reversion of G73S in the sequences with the double mutation only results in a shift of Δ*E* equivalent to 15% of the difference between the mean Δ*E*s in the ME and LE sequences. This suggests that while G73S certainly helps to entrench L90M, it is not required for the entrenchment of L90M and is not solely responsible for the entrenchment of L90M when present. Similar effects are observed for mutation I54V in the entrenchment of V82A and M46I and L90M in the entrenchment of I84V.

To uncover the clearest patterns of mutations that differentiate the LE sequences from the ME sequences, we performed principal component analysis (PCA) on the combined set of ME and LE sequences at PI-associated sites. The projections of the ME and LE sequences onto the first 3 principal components are shown in Figures 6 and S6. The first three principal components explain approximately 40% of the total variance when performed on the data corresponding to V82A, I84V, L90M (39.5%, 42.5%, 37.4% respectively). In the case of L90M, the first principal component clearly separates the most entrenched sequences from the least entrenched sequences while the second principal component separates variation within both groups. For V82A and I84V, a linear combination of the first two principal components separates the ME from the LE sequences, most likely due to variation between and within the most and least entrenching sequences being similarly large (which can be seen in the plots of hamming distance in Figure S6).

**FIG. 6:**
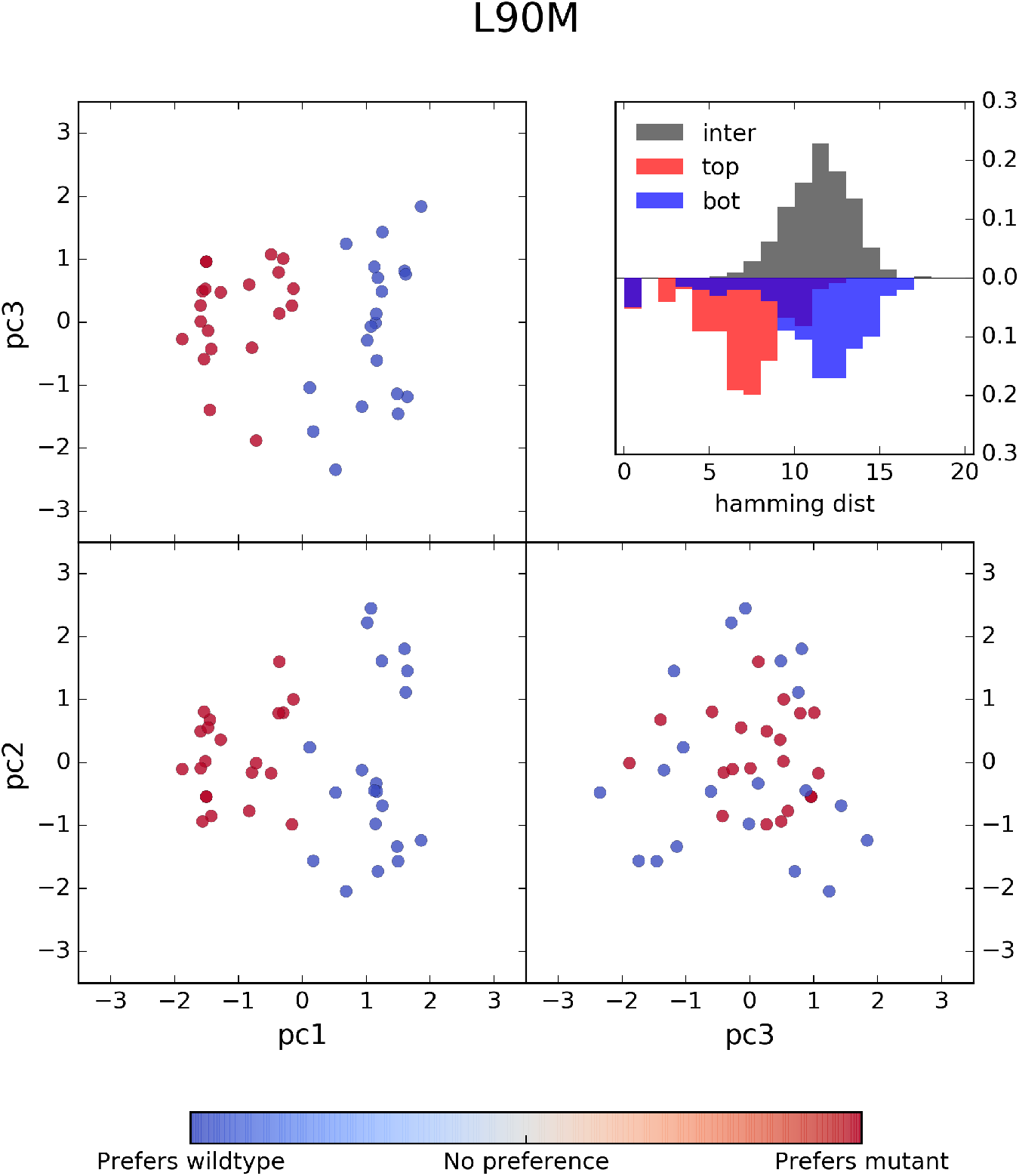
PCA analysis of most and least entrenching sequence backgrounds for primary resistance mutation L90M. Sequences from the 10^th^ and 90^th^ percentiles in Δ*E* of the sequences containing L90M and with a hamming distance of 10 from the consensus were labeled as “least entrenching” and “most entrenching”, respectively, and pooled. These sequences of length *L* = 93 encoded with a *Q* = 4 alphabet were transformed to bit vectors of length *LQ* and Principal component analysis (PCA) was performed on this set of transformed sequences. The projection of these sequences onto their first 3 principal compenents are shown above with the least entrenching sequences colored blue and most entrenching sequences colored red. The first principal component clearly separates the most from the least entrenching sequence backgrounds for L90M while the other two components explain variation within the two groups of sequences. Shown in the inset are the distributions of hamming distances between (gray) and within the most entrenching (red) and least entrenching (blue) sequences.

Examination of the first principal component (PC) eigenvector shows that the residues of at least 11 PI-associated sites contribute to the differentiation of the most entrenched (ME) sequences from the least entrenched (LE) sequences for primary mutation L90M, with residues K20F/I/V, M46I, G73S, V82V, and I84V contributing most strongly. Sequences from the two classes for which the first PC explains the most variation, measured as the hamming distance captured by the first PC, can be found in Table S1. Contributions from 11 sites is consistent with the average pairwise hamming distance of 11 between the most and least entrenched sequences, as seen in Figure 6 inset. Similarly, sets of 14 and 16 residues among the first two principal eigenvectors are responsible for the separation of ME and LE sequences for V82A and I84V, respectively (see Figure S6). These observations reinforce the point that while previously identified primary-accessory mutation pairs are important for acquisition and fixation of primary mutations, a model which captures epistatic effects collectively, like the Potts model, is needed to identify sequence backgrounds most likely to accommodate primary mutations.

Non-PI-associated polymorphisms also appear to modulate the entrenchment of primary resistance mutations, though the effect is secondary to that of PI-associated mutations. There exist sets of sequences, each with the same pattern of PI-associated mutations, that differ in entrenchment scores by as much as ΔΔ*E* ≈ 3, which corresponds to observable probabilities differing by more than an order of magnitude. This appears to be the result of strong positive and negative couplings that arise between non-PI-associated polymorphisms and certain PI-associated mutations. For example, we find that non-PI-associated mutations V11I, K43R/N, I66V, C67F/L/Q/E, I72V/L, T74A, P79A, and C95F all appear to regulate the entrenchment of L90M. Some of these residues lie in the hydrophobic core of the protease dimer, and subtle conformational changes in the hydrophobic core by these residues may play an important role in inhibitor binding (Mittal et al. 2012). A demonstration of this modulation is shown in Figure S7, where a common background sequence of 10 PI-associated mutations is shared by several observed sequences in the original MSA with varying number of additional polymorphisms. Two of these sequences are shown in Figure S7B, and contain one and six additional mutations respectively. Despite the complicated network of interactions, the presence of the additional five polymorphic mutations in the second sequence increases the entrenchment of L90M, with ΔΔ*E* = 2.39 when reverting L90M to L, which corresponds to ~ 10 fold increase in frequency.

These results present testable predictions, and we have included three pairs of sequences that we predict will be most and least entrenching for the primary mutations discussed here, which can be found in Table I. Using either replicative capacity or melting temperature as a proxy for fitness, it should be possible to verify experimentally whether the Potts model correctly predicts the relative frequencies upon reverting the primary mutation to wildtype for the selected sequences pairs listed in Table I.

**TABLE I:** Combinations of a most and least entrenching sequence corresponding to the entrenchment of the primary mutations V82A, I84V, and L90M.

| Position<sup>a,b</sup> | Consensus | V82A |  | I84V |  | L90M |  |
| --- | --- | --- | --- | --- | --- | --- | --- |
|  |  | ME<sup>c</sup> | LE<sup>d</sup> | ME | LE | ME | LE |
| **10** | L | I | I | F | L | I | L |
| 13 | I | I | V | I | V | I | I |
| **20** | K | R | K | K | R | I | K |
| **24** | L | I | L | L | L | L | L |
| **30** | D | D | D | D | N | D | D |
| **33** | L | L | F | F | L | L | L |
| 35 | E | D | E | E | D | D | E |
| **36** | M | I | M | M | I | M | I |
| 37 | N | N | N | D | S | S | D |
| 41 | R | K | R | R | R | R | K |
| **46** | M | L | M | I | M | I | M |
| **48** | G | G | G | G | G | G | V |
| **54** | I | V | V | V | I | I | V |
| 57 | R | R | R | R | K | R | R |
| **58** | Q | Q | Q | Q | E | Q | Q |
| **62** | I | I | I | I | V | V | V |
| **63** | L | P | P | P | P | P | P |
| 67 | C | C | F | C | C | C | C |
| **69** | H | H | H | H | H | H | Y |
| **71** | A | V | V | V | T | I | V |
| 72 | I | I | M | I | I | I | I |
| **73** | G | G | S | G | G | S | G |
| **74** | T | T | T | P | T | T | T |
| **77** | V | V | I | V | V | V | V |
| **82** | V | A | A | V | V | V | A |
| **84** | I | I | V | V | V | V | I |
| **88** | N | N | N | N | D | N | N |
| **90** | L | L | M | M | M | M | M |
| **93** | I | L | I | L | I | L | L |
| $\Delta\Delta E$ | | 6.93 | | 5.80 | | 5.52 | |
| Relative probability $ME/LE$ | | 1022 | | 330 | | 250 | |

<sup>a</sup> The residue at positions not listed is the subtype B consensus residue
<sup>b</sup> PI-associated positions are shown in bold
<sup>c</sup> most entrenching
<sup>d</sup> least entrenching

## III. DISCUSSION

The evolution of viruses under drug selective pressure induces mutations which are correlated due to constraints on structural stability and function that contribute to fitness. The correlations induce epistatic effects, a primary or accessory resistance mutation can be either stabilizing or destabilizing depending on the genetic background. Recently epistasis has become a focus for analysis in structural biology and genomics as researchers have begun to successfully link the coevolutionary information in collections of protein sequences with the structural and functional fitness of those proteins (Barton et al. 2016b, Butler et al. 2016, Ferguson et al. 2013, Figliuzzi et al. 2015, Hinkley et al. 2011, Hopf et al. 2015, Mann et al. 2014). In the current study, we have used the correlated mutations encoded in a multiple sequence alignment of drug-experienced HIV-1 protease sequences to parametrize a Potts model of sequence statistical energies that can be used as an estimator of stability and relative replicative capacity of individual protease sequences containing drug resistance mutations.

The most entrenching sequences are those at local fitness maxima, and accumulating mutations, as seen here as increasing hamming distance from the subtype B consensus sequence, unlocks pathways to these local fitness maxima (Gupta and Adami 2016). Up to 100–1000 times more probable than sequences that favor reversion to the consensus genotype, these highly resistant sequences observed in our MSA present a significant risk for the transmission of drug resistance to new hosts as they incur large fitness penalties for reversion. Indeed, we find that the entrenchment effect is strongest for L90M, which has been shown to revert very slowly in drug naive patients with transmitted drug resistance (Yang et al. 2015).

This work builds upon a large literature, ranging from experimental work (Chang and Torbett 2011, Henderson et al. 2012) and statistical analyses of covarying pairs of mutations (Rhee et al. 2007, Wu et al. 2003) to more advanced statistical models of patterns of mutations at many positions (such as Potts models) (Butler et al. 2016, Haq et al. 2009, 2012), to strengthen our understanding of the emergent properties of drug resistance in HIV-1 protease. We demonstrate that, while very important, the information conveyed by pairs of primary and accessory mutations only tells a small part of the story; the context of the full sequence background is really necessary to understand how primary resistance mutations become stabilized. The results presented here advance recent work in the field of using Potts models to study HIV evolution (Barton et al. 2016b, Butler et al. 2016) by providing systematic prospective predictions quantifying the influence of specific multi-residue patterns on the tolerance of drug resistance mutations.

Recent publications have reported that mutations near or distal to Gag cleavage sites play a role in promoting cleavage by drug-resistant and enzymatically deficient proteases, by selecting for mutations that increase substrate contacts with the protease active site, altering the flexibility of the cleavage site vicinity, or by as of yet unknown mechanisms (Breuer et al. 2011, Flynn et al. 2015, Fun et al. 2012, Kolli et al. 2009, Parry et al. 2011, Prabu-Jeyabalan et al. 2002). This suggests that viral coevolution of Gag with selective protease mutations may further stabilize multiple resistance mutations; thus, the analysis of protease mutation patterns can be extended to include amino acid substitutions within Gag and the Gag-Pol polyprotein. Furthermore, this type of analysis is not limited to protease and may be used to study the development of resistance in other HIV drug targets, such as reverse transcriptase and integrase, as well as other biological systems that develop resistance to antibiotic or antiviral therapies.

The Potts model is a powerful tool for interrogating protein fitness landscapes as it captures the correlated effects of many mutations collectively. The analysis presented here provides a framework to examine the structural and functional fitness of individual viral proteins under drug selection pressure. Elucidating how patterns of viral mutations accumulate and understanding their epistatic effects has the potential to impact design strategies for the next generation of c-ART inhibitors and therapies.

## IV. MATERIALS AND METHODS

### Sequence Data

Sequence information (as well as patient and reference information) was collected from the Stanford University HIV Drug Resistance Database (http://hivdb.stanford.edu) (Shafer 2006) using the Genotype-Rx Protease Downloadable Dataset (http://hivdb.stanford.edu/pages/geno-rx-datasets.html) that was last updated on 29/04/2013 (there now exists a more recent sequence alignment updated May 2015). There are 65,628 protease isolates from 59,982 persons in this dataset. From this dataset, 5,824 drug-experienced, subtype B, non-mixture, non/recombinant, and unambiguous sequences were extracted. Sequences with more than 1 gap and MSA columns with more than 1% gaps (positions 1–5 and 99) were removed, resulting in *N* = 5, 610 sequences of length *L* = 93.

For the comparison made in Figure S2, drug-naive subtype B non/mixture, non-/recombinant, and unambiguous sequences were extracted from the same downloadable dataset. As with drug/experienced sequences, gap-containing sequences and columns were removed, resulting in 13,350 sequences of length 89.

Mutations considered PI-associated were extracted from (Johnson et al. 2013) and https://hivdb.stanford.edu/dr-summary/resistance-notes/PI/: L10I/F/V/C/R, V11I, G16E, K20R/M/I/T/V, L24I, D30N, V32I, L33I/F/V, E34Q, M36I/L/V, K43T, M46I/L, I47V/A, G48V, I50L/V, F53L/Y, I54V/L/A/M/T/S, Q58E, D60E, I62V, L63P, I64L/M/V, H69K/R, A71V/I/T/L, G73S/A/C/T, T74P, L76V, V77I, V82A/F/T/S/L/I, N83D, I84V, I85V, N88D/S, L89I/M/V, L90M, I93L/M.

### Marginal Reweighting

Weights (*w*_*k*_) reciprocal to the number of sequences contributed by each patient were computed and assigned to each sequence. With these weights, estimates of the bivariate marginal probabilities were computed from the MSA of *N* sequences:

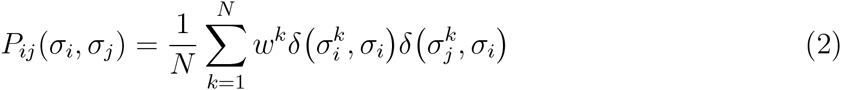

where 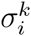 is the residue identity at position *i* of the *k*th sequence 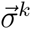, 0 < *w*^*k*^ ≤ 1 is the weight of sequence *k*, and delta *δ*(*α*, *β*) equals one if *α* = *β* and is otherwise zero.

Otherwise, all sequences are assumed independent; no reweighting was done to account for shared ancestry among these sequences. Phylogenetic trees of drug-naive and drugtreated HIV-infected patients have been show to exhibit star-like phylogenies (Gupta and Adami 2016, Keele et al. 2008), and thus phylogenetic corrections are not needed. Further, phylogenetic corrections based on pairwise sequence similarity cut-offs of 40% of sequence length or more as are common in studies utilizing direct coupling analysis (DCA) (Morcos et al. 2011, 2014,Weigt et al. 2009) of protein families would drastically reduce the number of effective sequences in our MSA and would lead to mischaracterization of the true underlying mutational landscape. Potts models of other HIV protein sequences under immune pressure have been parameterized with no phylogenetic corrections (Barton et al. 2016b, Ferguson et al. 2013, Mann et al. 2014).

### Alphabet Reduction

It has been shown that “reduced alphabets” consisting of 8 or 10 groupings of amino acids capture most of the information contained in the full 20 letter alphabet (Murphy et al. 2000). We expand on this notion by computing an alphabet reduction that has the least effect on the statistical properties of our MSA. In the context of model building, a reduced alphabet decreases the number of degrees of freedom to be modeled. This leads to a more efficient model inference (Barton et al. 2016a, Haldane et al. 2016).

Given the empirical bivariate marginal distribution for each pair of positions in the MSA using 21 amino acid characters (20 + 1 gap), the procedure begins by selecting a random position *i*. All possible alphabet reductions from 21 to 20 amino acid characters at position *i* are enumerated for every pair of positions *ij*, where *j* ≠ *i*, by summing the bivariate marginals corresponding to each of the 210 possible combinations of amino acid characters at position *i*. The reduction which minimizes the root square mean difference (RMSD) in mutual information (MI) content:

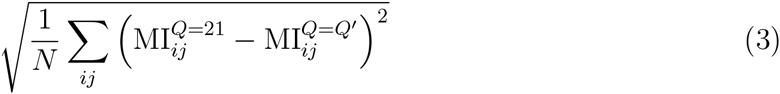

between all pairs of positions *ij* with the original alphabet size *Q* = 21 and reduced alphabet size *Q* = 20 is selected. The alphabet at each position i is reduced in this manner until all positions have position-specific alphabets of size *Q* = 20. This process is then repeated for each position by selecting the merger of characters which minimizes the RMSD in MI between all pairs of positions *ij* with the original alphabet size *Q* = 21 and reduced alphabet size *Q* = *Q*’, and is stopped once *Q* = 2.

Due to residue conservation at many loci in the HIV protease genome, the average number of characters per position is 2, and several previous studies of HIV have used a binary alphabet to extract meaningful information from sequences (Ferguson et al. 2013, Flynn et al. 2015, Shekhar et al. 2013, Wu et al. 2003). However, using a binary alphabet marginalizes potentially informative distinctions between amino acids at certain positions, especially PI-associated sites, that acquire multiple mutations from the wildtype. We found that an alphabet of 4 letters substantially reduces the sequence space to be explored during the model inference while providing the necessary discrimination between different types of mutant residues at each position. Additionally, the information lost in this reduction is minimal; Pearson’s *R*^2^ between the mutual information (MI) of the bivariate marginal distributions in 21 letters and in 4 letters is ≈0.995 (Figures S8, S9).

The original MSA was then re-encoded using the reduced per-position alphabet, and the bivariate marginals (Eq. 2) were recalculated using the reduced alphabet. Small pseudo-counts are added to the bivariate marginals, as described (Haldane et al. 2016). Briefly, instead of adding a small flat pseudocount such as 1/*N*, we add pseudocounts which correspond to a small per-position chance *μ* of mutating to a random residue such that the pseudocounted marginals *P*^pc^ are given by

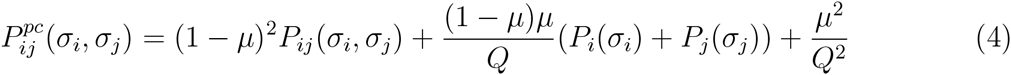

where we take *μ* ≈ 1/*N*.

### Maximum Entropy Model

Following (Mora and Bialek 2011), we seek to approximate the unknown empirical probability distribution 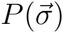 which describes HIV-1 protease sequences 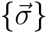 of length *L* where each residue is encoded in an alphabet of *Q* states by a model probability distribution 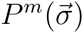. The model distribution we choose is the maximum entropy distribution, e.g. the distribution which maximizes

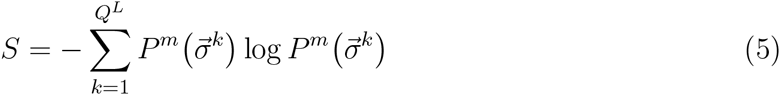

and has been derived by (Barton et al. 2016a, Ferguson et al. 2013, Mézard and Mora 2009, Morcos et al. 2011, Weigt et al. 2009) and others satisfying the following constraints:

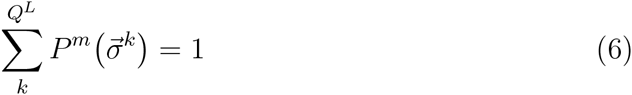

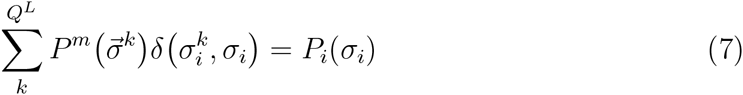

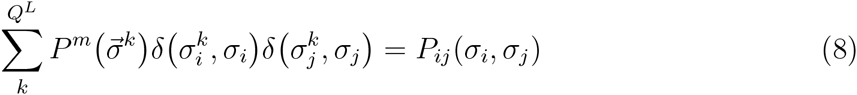

i.e. such that the empirical univariate and bivariate marginal probability distributions are preserved. Through a derivation using Lagrange multipliers not presented here (but can be found in (Ferguson et al. 2013, Mora and Bialek 2011)), the maximum entropy model takes the form of a Boltzmann distribution

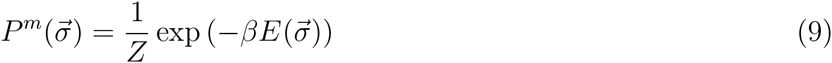

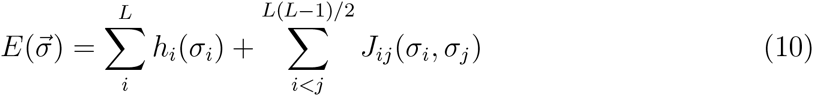

where the quantity 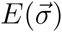 is the Potts Hamiltonian, which determines the statistical energy of a sequence 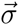, 1/*Z* is a normalization constant, and the inverse temperature *β* = 1/*k*_*B*_*T* is such that *k*_*b*_*T* = 1. This form of the Potts Hamiltonian consists of *Lq* field parameters *h*_*i*_ and 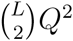 coupling parameters *J*_*ij*_ which describe the system’s preference for each amino acid character at site *i* and each amino acid character pair at sites *i*, *j*, respectively. In the way we present the Boltzmann distribution *P*^*m*^ ∝ exp (−*E*), negative fields and couplings signify favored amino acids preferences.

Not all the model parameters are independent. Due to the relationship between bivariate marginals *P*_*ij*_, *P*_*ik*_, *P*_*jk*_ and the fact that the univariate marginals can be derived entirely from the bivariate marginals, only 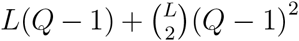 of these 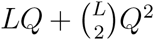 parameters are independent. Several schemes have been developed and used by others to fully constrain the Hamiltonian (see (Morcos et al. 2011, Weigt et al. 2009), for example). Further, the fully-constrained Potts Hamiltonian is “gauge invariant” such that the probablity 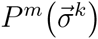 is unchanged by (a) a global bias added to the fields, *h*_*i*_(*σ*_*i*_) → *h*_*i*_(*σ*_*i*_) + *b*, (b) a per-site bias added to the fields *h*_*i*_(*σ*_*i*_) → *h*_*i*_(*σ*_*i*_) + *b*_*i*_, (c) rearrangement of field and coupling contributions such that *J*_*ij*_(*σ*_*i*_, *σ*_*j*_) → *J*_*ij*_(*σ*_*i*_, *σ*_*j*_) + *b*_*ij*_(*σ*_*j*_) and 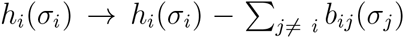, or (d) a combination thereof. Due to this gauge invariance, model parameters are over-specified and thus not unique until a fully-constrained gauge is specified, but the properties *P*^*m*^ and Δ*E*, among others, are gauge invariant and unique among fully-constrained gauges.

### Model Inference

Finding a suitable set of Potts parameters {*h*, *J*} fully determines the total probability distribution 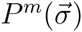 and is achieved by obtaining the set of fields and couplings which yield bivariate marginal estimates *P*^*m*^(*σ*_*i*_, *σ*_*j*_) that best reproduce the empirical bivariate marginals *P*^*obs*^(*σ*_*i*_, *σ*_*j*_). Previous studies have developed a number of techniques to do this (Balakrishnan et al. 2011, Barton et al. 2016a, Cocco and Monasson 2011, Ekeberg et al. 2013, Ferguson et al. 2013, Haq et al. 2012, Jones et al. 2012, Mézard and Mora 2009, Morcos et al. 2011, Weigt et al. 2009). Following (Ferguson et al. 2013), we estimate the bivariate marginals given a set of fields and couplings by generating sequences through Markov Chain Monte Carlo (MCMC) where the Metropolis criterion for a generated sequence is proportional to the exponentiated Potts Hamiltonian. The optimal set of parameters {*h*, *J*} are found through multidimensional Newton search, where bivariate marginal estimates are compared to the empirical distribution to determine descent steps. Unlike several inference methods referenced above, this method avoids making explicit approximations to the model probability distribution, though approximations are made in the computation of the Newton steps, and this method is limited by sampling error of the input emperical marginal distributions and by the need for the simulation to equilibrate. Also, the method is computationally intensive. A brief description of the method follows; see the supplemental information of Haldane et al. (Haldane et al. 2016) for a full description of the method.

Determining the schema for choosing the Newton step is crucial. In (Ferguson et al. 2013), a quasi-newton parameter update approach was developed, in which updates to *J*_*ij*_ and *h*_*i*_ are determined by inverting the system’s Jacobian, to minimize the difference between model-estimated and empirical marginals. To simplify and speed up this computation, we take advantage of the gauage invariance of the Potts Hamiltonian to infer a model in which *h*_*i*_ = 0 ∀ *i*, and we compute the expected change in the model marginals Δ *P*_*ij*_ (dropping the *m* superscript) due to a change in *J*_*ij*_ to first order by

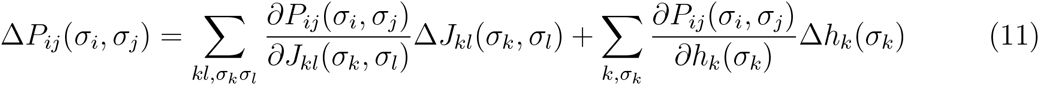

with a similar relation for Δ*P*_*i*_(*σ*_*i*_) The challenge is to compute the Jacobian 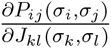 and invert the linear system in Equation 11, and solve for the changes Δ*J*_*ij*_ and Δ*h*_*i*_ given Δ*P*_*ij*_ which we choose as

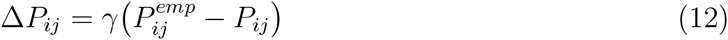

given a damping parameter *γ* chosen small enough for the linear (and other) approximations to hold.

The computational cost of fitting 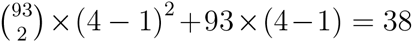, 781 model parameters on 2 NVIDIA K80 or 4 NVIDIA TitanX GPUs is approximately 4 hours. For a more thorough description of the inference methodology, see the supplementary information of Haldane et al. (Haldane et al. 2016).

### Experimental Comparison

Experimentally derived values for either melting temperature (*T*_*m*_) or viral infectivity via replicative capacity (RC) were mined from the results presented in (Chang and Torbett 2011, Henderson et al. 2012, Louis et al. 2011, Muzammil et al. 2003, van Maarseveen et al 2006). A csv file of the resulting mined data can be found in Supplementary Data 1.

## ACKNOWLEDGMENTS

This work was supported in part by National Institutes of Health (P50 GM103368 to W.F.F., B.E.T., R.M.L; R01 GM30580 to A.H., R.M.L.; S10 OD020095 to W.F.F., A.H., R.M.L.). We thank the supportive collaborative environment provided by the HIV Interaction and Viral Evolution (HIVE) Center at the Scripps Research Institute (http://hive.scripps.edu).

